# Cortexa – a comprehensive resource for studying gene expression and alternative splicing in the murine brain

**DOI:** 10.1101/2024.04.11.589045

**Authors:** Stephan Weißbach, Jonas Milkovits, Stefan Pastore, Martin Heine, Susanne Gerber, Hristo Todorov

**Affiliations:** Institute of Developmental Biology and Neurobiology (iDN), Johannes Gutenberg-University, 55128 Mainz, Germany; Institute of Human Genetics, University Medical Center, Johannes Gutenberg-University, 55131 Mainz, Germany; Institute of Pharmaceutical and Biomedical Sciences, Johannes Gutenberg-University Mainz, 55128 Mainz, Germany

## Abstract

**Motivation:** Gene expression and alternative splicing are strictly regulated processes that shape brain development and determine the cellular identity of differentiated neural cell populations. Despite the availability of multiple valuable datasets, many functional implications, especially those related to alternative splicing, remain poorly understood. Moreover, neuroscientists working primarily experimentally often lack the bioinformatics expertise required to process alternative splicing data and produce meaningful and interpretable results. Notably, re-analyzing publicly available datasets and integrating them with in-house data can provide substantial novel insights. However, such analyses necessitate developing harmonized data handling and processing pipelines which in turn requires considerable computational resources and in-depth bioinformatics expertise.

**Results:** Here, we present Cortexa – a comprehensive web-portal that incorporates RNA-sequencing datasets from the mouse cerebral cortex (longitudinal or cell-specific) and the hippocampus. Cortexa facilitates understandable visualization of the expression and alternative splicing patterns of individual genes. Our platform also provides SplicePCA – a tool that allows users to integrate their alternative splicing dataset and compare it to cell-specific or developmental neocortical splicing patterns. All gene expression and alternative splicing data have been processed in a standardized manner and they can also be downloaded for further in-depth down-stream analysis.

**Availability:** The data portal is available at https://cortexa-rna.com/

## 1 Introduction

Transcriptional regulation plays a crucial role in the developing and adult mammalian brain^1–3^. This includes especially gene expression^4,5^ and alternative splicing^6–9^ where major changes can be observed during development.

Among these factors, gene expression is the best-understood, and most extensively studied aspect. Particularly in neuroscience, it has been a major focus in the study of brain development^10^, cell identity^11,12^, and disease-related alterations^13–15^. However, a common issue with gene expression analysis when comparing results from heterogenous sources is the occurrence of batch effects that are unrelated to biological factors, due to differences in sequencing technology, experimental handling, and bioinformatic processing^16–20^.

Alternative splicing (AS) of precursor mRNA is a fundamental process that enables the generation of various transcripts and subsequent proteins from the same gene. Therefore, it significantly increases the available transcriptomic variability^21^. AS is particularly important in the central nervous system, playing a vital role during cortical development^6,21^ and in the determination and maintenance of neuronal cell identity^22^. Moreover, AS is an important regulator of gene expression since it can initiate the degradation of mRNA by introducing premature stop codons which leads to non-sense-mediated decay^23^.

Although several high-quality datasets focusing on AS in the murine brain are freely available, interpreting the different types of splicing events for individual genes remains challenging. Moreover, harnessing the potential of multiple datasets requires a harmonized processing strategy to ensure the comparability of results. A limitation of existing data portals, namely, the single cell atlas of the Allen Brain Institute^24^, Neuron Subtype Transcriptome^25^, or Brain RNA-Seq^26^, is that they are focused on gene expression and do not offer the analysis of custom data in the context of cell-specific or developmental changes in alternative splicing.

Here, we introduce Cortexa – a novel data portal for accessing a variety of high-quality neocortical and hippocampal transcriptomic datasets, analyzed for gene expression and alternative splicing. Batch effects between different studies have been minimized using a standardized analysis pipeline (a). We offer easily interpretable summaries and visualization of results that allow a broad range of scientists to explore the expression and AS patterns of individual genes. Additionally, we developed SplicePCA – a tool that performs a principal component analysis of splicing events for a selected gene set and enables the investigation of splicing patterns related to developmental changes or variations across cell types^6,27–29^. All standardized datasets included in Cortexa are publicly available for download, allowing users to integrate them into their research easily.

## 2 Methods

### 3.1 Datasets

We analyzed publicly available *in vivo* paired-end RNA sequencing data of the mouse cerebral cortex and hippocampus (d) with a minimum read length of 100bp (mRNA). We downloaded the sequencing data from NCBI SRA or GEO respectively. Specifically, we used SRP055008^6^, GSE133291^22^, and GSE96950^9^. Further datasets can easily be integrated, refer to https://cortexa-rna.com/datasets.

### 3.2 RNA-seq analysis

We used a standardized RNA-sequencing pipeline (a) to analyze the transcriptomic data for gene expression and alternative splicing. In brief, we trimmed the reads for adapter sequences with BBDuk (version 39.01)^30^. The trimmed reads were aligned to the reference genome mm39 (released 19.10.2022) downloaded from Gencode using STAR (version 2.7.10b)^31^. FeatureCounts, provided by SubRead (version 2.0.6) ^32^, was used to quantify the expression of each respective gene. All gene expression counts were normalized to transcripts per million (TPM).

We utilized rMATS turbo (version 4.1.2)^33^ with default settings to detect AS events. We analyzed the data for five alternative splicing events, cassette exon (skipping exon), mutually exclusive exons, intron retention, alternative A5’ splice site, and alternative A3’ splice site (b), as defined in rMATS^33^. The coverage for sashimi plots was analyzed with bamCoverage (version 3.5.2) of deepTools2^34^ converted to wig with Encode bigWigToWig.

### 3.3 Webapp

The web application was built with Next.js frontend framework, utilizing SQLite database for backend data storage. TypeScript enhances code maintainability and type safety, while Tailwind CSS streamlined styling. Prisma serves as the ORM tool for efficient database management. The application follows REST principles for communication between frontend and backend components, optimizing interoperability and scalability. The website is hosted on servers of the Johannes Gutenberg University, Mainz, Germany. A detailed tutorial on how to interpret alternative splicing events presented in Cortexa is provided at https://cortexa-rna.com/tutorial.

### 3.4 Visualization of Genes

Gene expression was normalized as transcripts per million (TPM) and represented as a heatmap for each dataset. Alternative splicing events (b) are visualized as sashimi plots.

### 3.5 SplicePCA

The principal component analysis (PCA) is performed on the averaged percentage spliced-in values of each experimental group. Only splicing events without missing values for any selected sample are considered for further analysis. It is possible to select specific genes and alternative splicing events, respectively, or to use all events at once. Moreover, SplicePCA allows users to integrate their in-house analyzed output files (rMATS). We recommend processing the files as described in section 3.2 and in the tutorial https://github.com/s-weissbach/cortexa_SplicePCA_example/.

### 3.6 Example Usage of SplicePCA

To validate SplicePCA, we obtained *Nova2*-knock-out (KO) and wild-type (WT) data from NCBI GEO with the accession number GSE103314^35^. We performed quality control, trimming, alignment, and alternative splicing analysis as described in section 3.2 RNA-seq analysis. Subsequently, the cassette exons from the rMATS output file were up-loaded to https://cortexa-rna.com/pca and analyzed in the context of developmental^6^ and NPC/neuron-specific^9^ alternative splicing events. Next, the results from SplicePCA were downloaded and plotted using matplotlib.

## 3 Application and discussion

Alternative splicing is a prevalent regulatory mechanism in the brain that plays an important role during development and in specifying and maintaining neural cell types^6,7,9,22,28,35,36^. However, *Mus musculus* has ∼22,000 protein-coding genes^37^ of which almost all multi-exon genes undergo alternative splicing^38^. Functional implications of these alternative splicing events remain in many cases elusive. Cortexa is an easy-to-use web tool to access alternative splicing events for genes of interest in a developmental and neuronal cell-type-specific context to formulate and investigate research hypotheses.

Additionally, principal component analysis has proven to be a powerful tool to summarize alterations in the alternative splicing landscape ^6,27– 29^. Using SplicePCA, researchers can select splice events for specific genes, integrate their data, and interpret their results in the context of cortical development and specific neuronal cell types. To showcase the usefulness of this approach, we re-analyzed cortical *Nova2*-KO and WT samples at the age of E18.5^35^ and used the SplicePCA tool. NOVA2 belongs to the class of RNA-binding proteins, governing alternative splicing during cortical development and in mature neurons^39^. Specifically, NOVA2 is required to regulate neuronal migration through splicing *Dab1* which is part of the Reelin pathway^2^. By using SplicePCA, we revealed a striking effect of *Nova2*-KO during cortical development. E18.5 knockout samples were located between E14.5 and E16.5 wild-type samples on the developmental splicing trajectory, indicating a less mature splicing pattern (c). Thus, NOVA2 splicing activity contributes significantly to the splicing changes between E14.5 to E18.5, as it was reported previously^2,35,40^. These results support the relevance of SplicePCA which combines available datasets with new datasets.

Cortexa thus allows the use of publicly available data without extensive re-analysis, which would require significant computational resources.

## 4 Conclusion

In summary, Cortexa gives access to high-quality, publicly available transcriptomic data to a broad range of scientists without the need to gain expertise in the computational aspects of gene expression and alternative splicing analysis. For experienced users, SplicePCA offers a powerful tool to summarize alterations in the alternative splicing landscapes upon experimental manipulation and compare results to the normal splicing trajectory during cerebral cortical development or in distinct neuronal cell types. Ultimately, Cortexa offers an easy-to-use extensive platform for gaining novel insights into gene expression and alternative splicing regulatory processes of the mouse brain.

## Author contributions

SW and HT conceived the idea and conceptualized the work. HT, SG, and MH supervised the project. SW and SP performed the bioinformatic analysis. JM and SW implemented and tested the website. SW created visualizations. SW and HT wrote the manuscript. MH and SG acquired funding and edited the manuscript.

## Data Availability

The *Nova2*-KO data that were used to test SplicePCA and a tutorial are available on GitHub at https://github.com/s-weissbach/cortexa_SplicePCA_example/.

---

**Figure 1.**
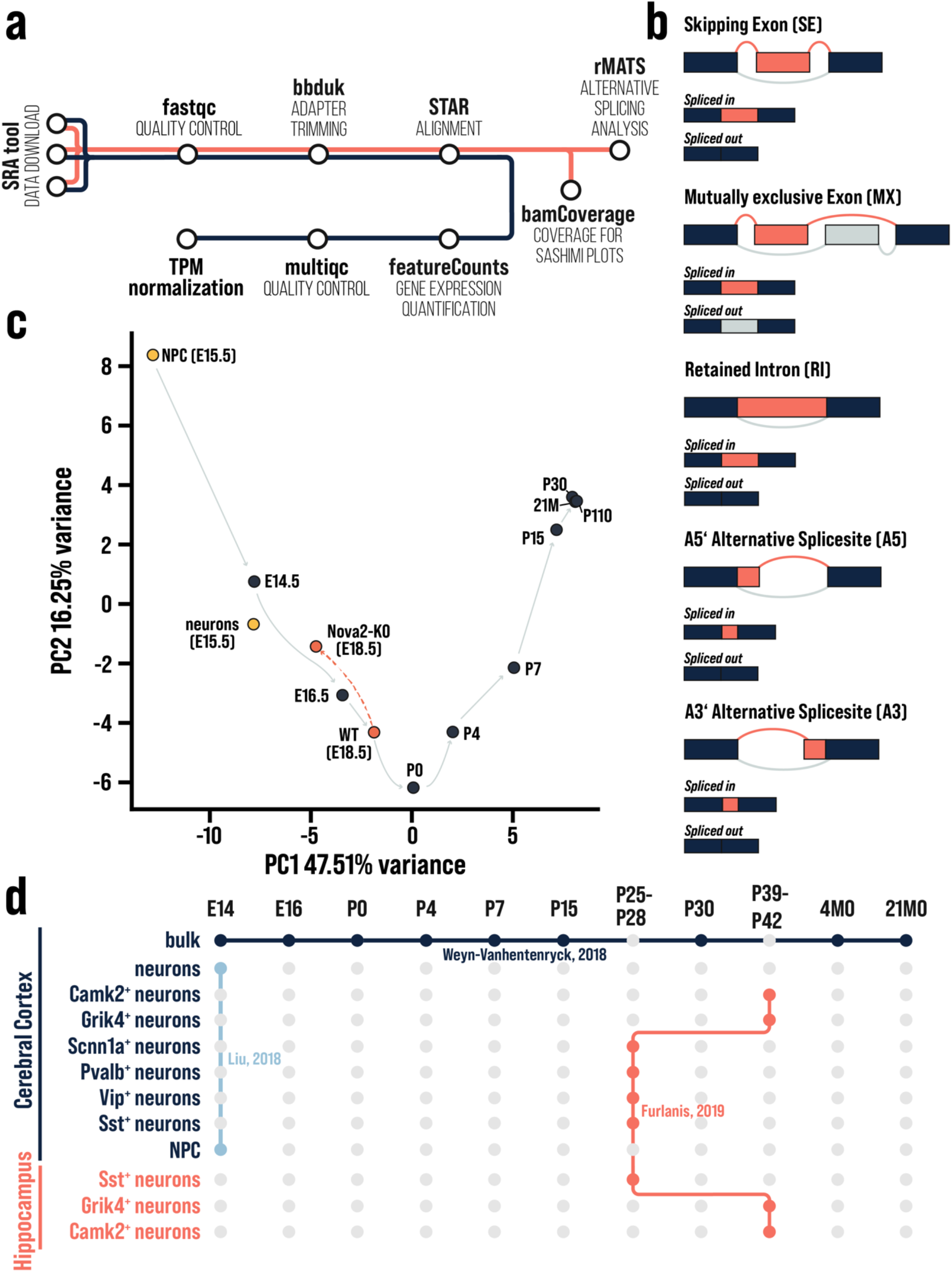
a) Standardized RNA-seq pipeline used for gene expression (blue) and alternative splicing (orange). b) Visualization of possible alternative splicing events and the splicing outcome referred to as spliced-in and spliced-out. c) Cassette exon data of cortical samples of WT (E18.5) and Nova2-KO (E18.5), analyzed with SplicePCA. In control conditions, PCA of alternative splicing data forms a characteristic bell-shaped trajectory with P0 as its inflection point. The conditional knock-out of Nova2 resulted in a substantial deviation from the normal splicing trajectory. According to the PCA, the Nova2-KO samples (E18.5) were associated with a less mature splicing pattern than E16.5 wild-type samples. d) Publicly available datasets integrated into Cortexa: SRP055008^6^, GSE133291^22^, and GSE96950^9^

